# ChIPSummitDB: A ChIP-seq-based database of human transcription factor binding sites and the topological arrangements of the proteins bound to them

**DOI:** 10.1101/652420

**Authors:** Erik Czipa, Mátyás Schiller, Tibor Nagy, Levente Kontra, László Steiner, Júlia Koller, Orsolya Pálné-Szén, Endre Barta

## Abstract

ChIP-Seq reveals genomic regions where proteins, e.g. transcription factors (TFs) interact with DNA. A substantial fraction of these regions, however, do not contain the cognate binding site for the TF of interest. This phenomenon might be explained by protein-protein interactions and co-precipitation of interacting gene regulatory elements. We uniformly processed 3,727 human ChIP-Seq data sets and determined the cistrome of 292 TFs, as well as the distances between the TF binding motif centers and the ChIP-Seq peak summits.

ChIPSummitDB enables the analysis of ChIP-Seq data using multiple approaches. The 292 cistromes and corresponding ChIP-Seq peak sets can be browsed in GenomeView. Overlapping SNPs can be inspected in dbSNPView. Most importantly, the MotifView and PairShiftView pages show the average distance between motif centers and overlapping ChIP-Seq peak summits and distance distributions thereof, respectively.

In addition to providing a comprehensive human TF binding site collection, the ChIPSummitDB database and web interface allows for the examination of the topological arrangement of TF complexes genome-wide. ChIPSummitDB is freely accessible at http://summit.med.unideb.hu/summitdb/. The database will be regularly updated and extended with the newly available human and mouse ChIP-Seq data sets.

## INTRODUCTION

ChIP-seq (<u>Ch</u>romatin <u>I</u>mmuno<u>P</u>recipitation followed by high-throughput sequencing) is a powerful technique, which reveals the genome-wide positions of those DNA sequences that co-precipitate with a given protein, which was used to generate the antibody for the IP, (1,2). The interaction between the protein and the DNA can be direct or indirect. Direct interactions can be specific, i.e. when a protein (transcription factor) recognizes and binds to a DNA sequence motif; or it can be nonspecific, as in the case of histones or cohesins (3–5). Indirect interactions between DNA and proteins occur through transcriptional regulatory complexes and/or DNA looping. In such cases, the cognate binding site for the given TF is not present under the ChIP-seq peaks (Additional file 1) (6).

In a typical primary ChIP-seq analysis pipeline the sequence reads are mapped to a reference genome, areas with the highest coverage (peaks) are determined, and the enriched *de novo* or known motifs at the peaks are identified. These steps are followed by downstream analyses, which typically involve peak annotation, comparison of different ChIP-seq experiments, and visualization, for example generating profiles, heat-maps and Venn-diagrams) (7). The most critical step in such a pipeline is the peak calling. Different peak calling algorithms provides different results and the number of the determined peaks also depends on the number of the sequenced reads (8).

Today, raw data from more than 85000 human and mouse ChIP-seq experiments are available (9), which gives the opportunity to perform further analyses and/or to set up secondary databases using those data. Previously, such databases have been built based on different parameters of ChIP-seq analyses. Some databases (CODEX, BloodChIP, hmChIP) put more focus on the experimental metadata collection and the classification of the experiments by the cell type (10–12). In addition, CODEX provides a visualization tool for examining peaks (10). Other databases, for example Cistrome Data Browser, GTDR, ChIP-Atlas, Factorbook, carry out different downstream analyses to show further details (13–16). Most of these databases are not only a simple collection of ChIP-seq data and a display of ChIP-seq peaks. Factorbook, for example, has an interactive tool to examine the nucleosome and histone modification profiles around the ENCODE TF ChIP-seq peaks (16,17). The gene transcription regulation database (GTRD) project, among other things, focuses on improving the peak calling procedure (14). They use several peak calling algorithms and make clusters of overlapping results. ChIP-Atlas provides a tool for extensive co-localization and enrichment analyses (15). TFBSbank focuses on annotating genomic localizations, finding co-binding proteins, and searching for *de novo* and known motifs within the peaks (18). The Cistrome Data Browser combines ChIP-seq data with chromatin accessibility data and provides a convenient web interface to browse and download these data (13). Most of the above-mentioned databases contain not only human but mouse data too (Cistrome Data Browser, Factorbook, CODEX, hmChIP), and, in some cases, ChIP-seq data for other species (GTRD, ChIP-Atlas, TFBSbank).

Enhancers are distant regulatory elements relative to transcription start sites (19). They can be characterized by TF binding (GTRD, TFBSbank), certain histone marks (SEdb), and enhancer transcription (HACER) (20,21). Because both TF binding and histone modifications can be detected by ChIP-seq, secondary ChIP-seq database tools can predict enhancer sequences as well if they determine the cognate binding sites under the peaks of TF ChIP-seq experiments. The ChIP-seq peaks and the protein-binding DNA-motifs under them can be further analyzed, and the resulted data can be integrated into a higher level database, such as the TRRUST (22).

ChIP-seq databases can provide tools to search, analyze, visualize, and download existing ChIP-seq data. Our previous results, however, demonstrated that ChIP-seq peak summits can also help to understand the topological arrangements, the spatial position(s) of different proteins bound to the DNA double helix. Therefore, our aim was to extend our CCCTC-binding factor (CTCF) peak summit-based analysis to every available ChIP-seq transcription factor, which was examined in human ChIP-seq experiments (23). For the study, we have manually selected 3727 ChIP-seq experiments, which representing a very large number of human TFs and co-factors (9). Since determining the correct positions of peak summits is critical for the analysis, we have developed a robust peak filtering pipeline, by which the positions of peak summits and the mapped TFBS motifs could be compared. Therefore, based on the ChIP-seq peak regions for each TF, we defined the corresponding consensus motif binding site sets for each of them. In addition to the consensus motif binding site sets, the ChIPSummitDB contains the distances between each pair of mapped consensus motifs and ChIP-seq peak summit.

The web interface of the ChIPSummitDB can display data in different views. Using the GenomeView module, users can browse the genome for ChIP-seq peaks and consensus motif binding site sets. The MotifView option can display the average distances between the centers of consensus motif binding sites and the ChIP-seq peak summit positions for each ChIP-seq experiment with overlapping peaks. Three different ChIP-seq experiments can be compared in the PairShiftView module. Using the dbSNPView option, users can evaluate whether a dbSNP entry overlaps with one or more consensus motif binding sites from our database (24).

In addition to browsing all processed data, the analysis of the scatterplots provided in the MotifView led us to hypothesize that the extent of the standard deviation of the motif center vs peak summit distances may be proportional to the closeness of the given protein to the DNA double helix.

## MATERIAL AND METHODS

To construct the ChIPSummitDB, primary raw read data of ChIP-seq experiments and the accompanying metadata were obtained from the NCBI SRA database (9). Processing of the downloaded data into their final appearance in ChIPSummitDB is summarized in Fig. 1. To determine peak regions, peak summits, consensus motif positions, and the distances between peak summits and motif centers, we carried out eight different processing steps (Additional file 1: Figure S1-S7). The scripts used during the process were deposited to the GitHub repository (https://github.com/summitdb). Below we provide some description of each step while more details and explanation of them are on the ChIPSummitDB web site (http://summit.med.unideb.hu/summitdb/Documentation.php) and in Additional file 1.

**Fig. 1.**
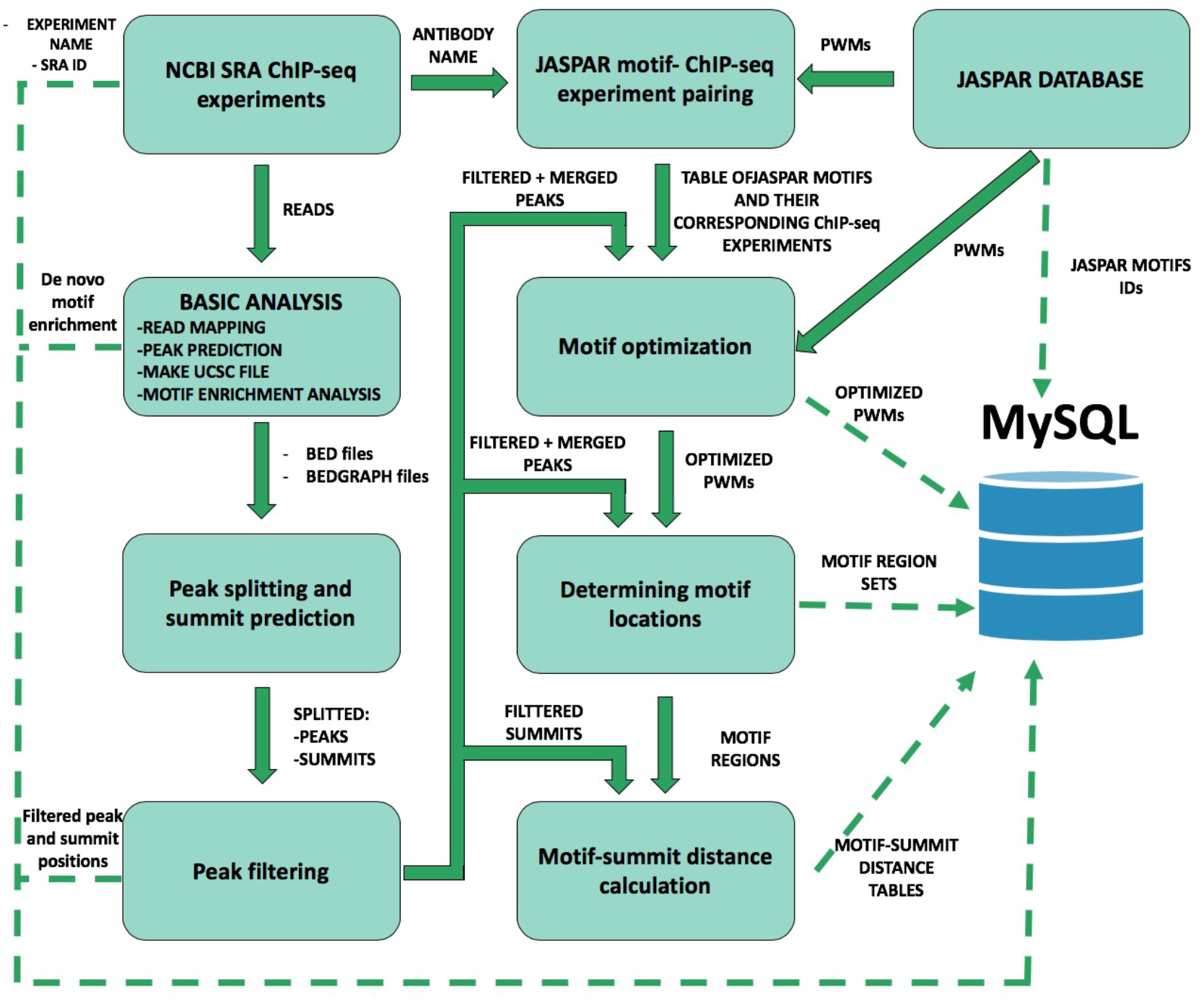
Schematic overview of ChIP-seq data processing and imported content from MySQL. The analysis steps and data conversion are marked with thick arrows. The uploaded results/files are represented with dashed lines. A vast majority of processed data is available on ChIPSummitDB, including the predicted peak regions, optimized JASPAR CORE PWMs, identified TFBSs, and calculated protein position information.

### 1. ChIP-seq data collection from public databases

We searched for human ChIP-seq experiments in the NCBI SRA database according to its status as per November 1, 2017. We used the NCBI’s run selector feature to download all available metadata associated with the selected experiments (9). A custom PERL script was used to process the downloaded data in XML table format and to give a unique (descriptive) name to every experiment. The names include the species, the tissue, the cell line (if available), the pathology (eg. normal or cancer), the ChIP target protein, and the experiment’s SRA database ID. Our aim was to restrict the analysis to transcription factors and to other non-histone proteins, thus experiments with other type of proteins were filtered out by using a script. For simplicity and to avoid redundancy, we processed only the normal (without any specific treatment, which influence the TF-DNA interaction) ChIP-seq experiments. The final list contained 4052 experiments and was converted into a table with the proper format (BED, BEDGRAPH) for further processing (Additional file 1: Figure S1).

### 2. Basic ChIP-seq analysis

For the basic processing, we used a modified version of our ChIP-seq_anal BASH script (Additional file 1: Figure S1) (7). Briefly, the script needs two input files. The first is the above-mentioned table with the experiments, while the second contained the SRR IDs for each SRX ID. After downloading the SRR files and converting them into fastq format, we mapped the reads to the hg19 (GRCh37) human reference genome using the Burrows-Wheeler Aligner (BWA) program (25,26). Peak calling, generation of the bedgraph coverage files, and *de novo* identification of protein-binding DNA motifs were performed by using the HOMER package. The output of the analysis contained the BAM files with the read sequences, the HOMER tag directories, the peaks in BED file format, the BEDGRAPH files, and both the *de novo* identified and previously known motifs, in a single html report (27).

### 3. Peak splitting and summit prediction

The basic analysis provided the peak regions for each ChIP-seq experiment in two forms, a BED file containing the borders of the HOMER predicted peak regions and a BEDGRAPH file with the coverage values of the extended reads within these peak regions (27). It is possible to have more than one binding site within a peak region, which can result in more than one summit. We employed the PeakSplitter program to determine such summit positions using the BEDGRAPH files as input (28). The result of this step is a BED file for each experiment containing the positions of the identified summit(s) for each peak (Additional file 1: Figure S2).

### 4. Peak filtering

The usefulness of the ChIPSummitDB largely relies on the correct determination of peak regions. Different peak -finding algorithms can give surprisingly diverse peak sets using the same ChIP-seq experiments. A number of parameters in the experiment, such as the number of the cells in the biological sample, the conditions of the sonication, the quality of the antibody, the library preparation, and especially the depth of the sequencing affect the number of detected peaks. There are different approaches to determine the biologically most relevant peak regions. One approach was to apply different peak calling programs and find the consensus peak sets from the results. Furthermore, we also developed a different approach, in which we used only the HOMER peak calling program, but applied a rigorous filtering, which was based on the shape of the peaks, to reduce the number of false positives/artifact peaks in the peak sets obtained in the previous step for downstream summit based topology analysis (Additional file 1: Figure S3). Using this approach, we filtered out about 35 percent of all peaks (27) (Additional file 3).

### 5. Assigning consensus motifs to TF ChIP-seq experiments

We assume that if the antibody used during the immunoprecipitation is against a TF, then the cognate binding site for that TF will be enriched in the peak set of the given ChIP-seq experiment. During the basic analysis, we determined the enriched motifs for each peak set. The problem was that even if the immunoprecipitated TF was the same, the resulting *de novo* motif could be slightly different. Also, the *de novo* motif finding algorithms usually give more than one enriched motif and it is unclear whether the best one is the cognate binding site. To precisely determine the peak summit motif center distances, however, we need to use the same binding site for every experiment with the same TFBS. Therefore, we chose a reverse approach. Based on the antibody used during the immunoprecipitation, we assigned a JASPAR core consensus motif to each experiment by hand. This resulted in a table where there is a corresponding JASPAR consensus motif for each TF experiment (Additional file 1: Figure S4, Additional file 2) (29).

### 6. Motif optimization

Many of the consensus motifs in the JASPAR database are based on *de novo* motif finding in ChIP-seq peak regions (29). Since we now have a good collection of representative ChIP-seq experiments for the targeted transcription factors, we decided to further optimize the consensus motif matrices. For this, we first merged the peaks of the experiments belonging to the same consensus motif, and then we used the HOMER package to optimize the matrices on these merge peak regions (Additional file 1: Figure S5) (27). The resulting optimized matrices were further inspected and adjusted manually. The motif optimization resulted in more than one similar motif in a few cases. The decision between them couldn’t be automatized. In these cases, the most analogous motif was chosen.

### 7. Remapping motifs

Once we had all the optimized consensus motif matrices, we needed to map them back to the genome. For each optimized JASPAR consensus motif matrix, we had a list of corresponding ChIP-seq experiments (Additional file 1: Figure S4) (29). We took the peak regions for those experiments and merged them to create the merged peak regions for each matrix. This is an important and, so far, a unique step among the similar ChIP-seq databases. Using this method, we specifically determined the possible transcription factor binding sites. For mapping, we used three different programs and we kept only the positions where at least two of them gave a hit for the final consensus motif binding site sets (Additional file 1: Figure S6) (27,30,31).

### 8. Motif center and summit distance calculation

After the peak filtering in step 4, we got the list of peak summits for each ChIP-seq experiment. In step 7, we got the ChIP-seq verified positions of TFBSs for each consensus motif. The majority of the peak regions do not contain the cognate binding site, even for TF ChIP-seq experiments. We hypothesize that in these cases the given TF can be part of a complex, which is bound to the DNA through another TF, which in turn is bound to its cognate binding site, which is also present in our consensus binding site set. Therefore, we investigated such cases as well. To do this, we calculated every distance between consensus motif centers and the nearby ChIP-seq peak summit positions. The calculation is motif based, which means that the output is a list for each consensus motif set (i.e. a given transcription factor binding site), which contains the distances of every peak summit inside the 50 bp range of the given consensus motif binding site (Additional file 1: Figure S7) (32). Practically, if we see a consensus motif (e.g. CTCF), we will have the ChIP-seq verified instances of that motif in the genome (88906) and a list of distances for each of the 3727 ChIP-seq experiments. These lists were processed further in the database. The number of distances in these lists were used as a cutoff value in the MotifView. The standard deviation of the distances in these lists was also calculated and indicated in the Y-axis of the MotifView.

This processing of raw ChIP-seq read and metadata resulted in the following tables and files:

1. The HOMER *homerpeaks.txt files for each experiment. These files contain the peak regions and other parameters used in peak finding.
2. A peak region and a summit bed format file for each experiment.
3. A table with all the metadata related to the ChIP-seq experiments.
4. A table with the average distances between each consensus binding site set and experiment.
5. A table with all the distances between individual consensus binding sites and each overlapping peak summit.
6. The HOMER *de novo* motif finding results for every experiment.
7. The optimized consensus motif matrices for every consensus binding site set.

Based on these files and tables, the physical and logical scheme of the database was developed using Oracle SQL Developer Database Modeler (version 4.2). For data storage, MariaDB (version 5.5.60) was chosen because it is free and open source. The website is hosted in CentOS Linux with Apache HTTP server and pages were developed using PHP. Graphs and other visual elements representing statistical results were made using the D3.js (version 3) JavaScript library.

## RESULTS AND DISCUSSION

The ChIPSummitDB addresses two main aims. First, the database is a comprehensive collection of ChIP-seq verified binding sites for 292 different TFs. Second, the database provides a new tool for analyzing the topological relationship between bound and co-bound proteins on the TFBSs.

There are six different views for exploring the database. From the main page, the user can either choose one of them under the search view tab or click any of the example pictures in order to go directly to a view page.

### User interfaces

**MotifView** is the summary page for each of the examined TFs. The scatterplot presented in MotifView shows the average distances of the overlapping peak summits relative to the center of the adjusted TFBS. In the Y-axis, one can choose between showing the number of overlapping peak summits and TFBS pairs or the standard deviation of those distances.

Each point on the plot represents one ChIP-seq experiment. Circles represent experiments where the antibody was against a TF with a defined TFBS, while triangles show co-factors and other proteins attached to the primarily bound TF. On the top of the scatterplot box there are three buttons. On default, all the experiments are displayed which have the element number above the threshold set at the left side bellow the box. In many cases when the “Minimum overlap number between motifs and peaks of experiment” threshold is set to too low, the box is overloaded with scatters. In this case there are two options. The “Only direct binding” button selects only those ChIP-seq experiments, which have been done with the antibody corresponding to the given consensus motif. The second option is to hide all the scatters. It allows then to individually change the experiments to be shown because on the right side of the plot, there is a color-coded list of antibodies or cell types used in these experiments. The list can be sorted alphabetically or by the number of occurrences. By clicking on these squares (TFs) or triangles (co-factors and other proteins), one can change the displayed experiments. For example, the user can choose and examine only the experiments belonged to one or more antibodies or cell types. This allows users to flexibly change the displayed experiments. On the scatterplot, one can select up to three experiments for further analysis in the other views.

The **PairShiftView** allows users to examine the distribution of individual peaks vs. summit consensus motif distances for three experiments. To smooth the graph, a 5 bp rolling bin is used in the histogram. Each curve represents one ChIP-seq experiment. During the creation of the database, both the summit locations for all peaks for a given experiment and the consensus motif binding sites of the chosen TF have been determined. In the latter case, only those binding sites that have been counted are shown, where the binding site in the genome overlaps at least one peak of a ChIP-seq experiment immunoprecipitated with the antibody against the corresponding TF. In the histogram, the highest points of the curves show the most likely topological position for a given protein on the DNA double helix relative to the consensus motif. The height of the curve depends on the number of overlapping consensus motif summit pairs (element number). In most cases, the curve has one peak, but sometimes it also has a shoulder. This indicates a more complex topological arrangement or a consequence of other, still unknown, reasons. It is remarkable for example that in many cases the distance between the main summit and the shoulder (or between two summits from different experiments with the same antibody) is approximately 11 or 22 bp (Fig. 2) (http://summit.med.unideb.hu/summitdb/paired_shift_view.php?exp1=419&exp2=1960&exp3=3681&motive=FOXA1&motifid=77&limit=25&low_limit=-25&formminid=1&formmaxid=10000&mnelem=100&formmaxelem=120000), which represents one or two turn in the double helix and can be interpreted as a protein on the same side of the DNA (33).

**Fig. 2.**
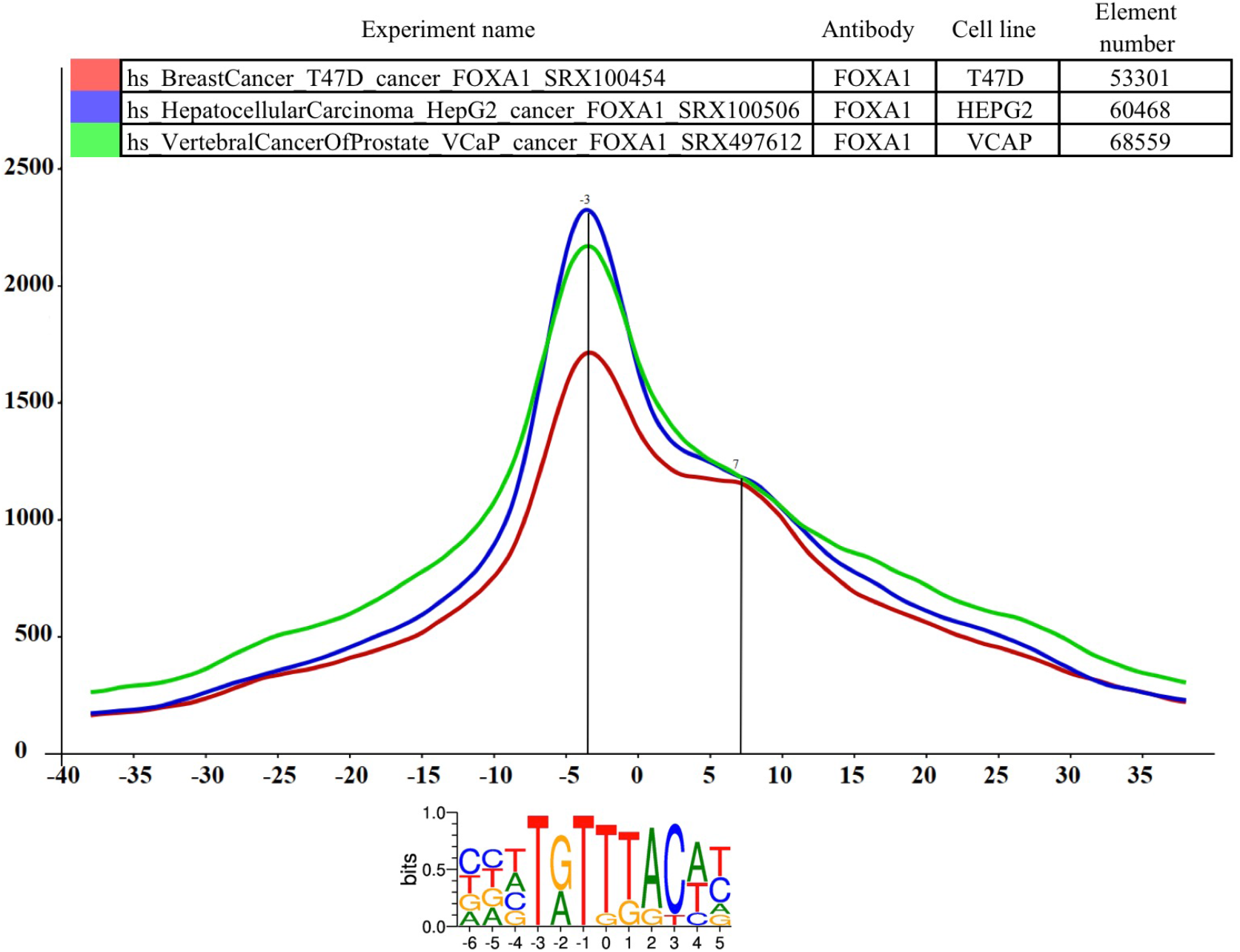
The distance distribution of FOXA1 summits relative to the motif centers of FOXA1 binding sites. The horizontal axis represents the distance of summits in different cell lines (T47D (SRA ID: SRX100454, red curve), HepG2 (SRA ID: SRX100506 blue curve) and VCaP (SRA ID: SRX497612, green curve)) relative to the FOXA1 motif center. The vertical axis represents the distance frequencies. A rolling mean with a 5 bp window was applied to smooth the frequency curves. The distance between the maxima (main summit, maxima at −3 bp) and the shoulder (7 bp) is approximately 10 bp. Element numbers in the table indicates the number of peak regions obtained in a ChIP-seq experiment, which overlap with a particular consensus motif binding site set. Figure is adapted from ChIPSummitDB website: http://summit.med.unideb.hu/summitdb/paired_shift_view.php?exp1=419&exp2=1960&exp3=3681&motive=FOXA1&motifid=77&limit=25&low_limit=-25&formminid=1&formmaxid=10000&mnelem=100&formmaxelem=120000

Although the highest point of the curve in the PairShiftView and the position of the corresponding scatter in the X-axis indicate the same topological positioning of the given protein on the DNA double helix, these values can be different. In most cases, this is because in the histogram the distribution curve either has a shoulder or is simply asymmetric. Therefore, we assume that while the scatterplot in the MotifView gives a good overview of the differences in the topological arrangements of the given consensus motif experiments pairs, the PairShiftView is a more accurate tool to examine the relative topological arrangements of the three chosen experiments.

Although in most cases one gets to the PairShiftView from the MotifView by selecting three experiments, the user can change both the consensus motif and the selected experiments from inside the PairShiftView. This allows a quick comparison of any three experiments.

Besides seeing the topological arrangement of three chosen immunoprecipitated proteins relative to the given motif center, the extent to which the peaks of the three ChIP-seq experiments share the same binding sites is also interesting. The **VennView** is designed to allow this. During the processing of the ChIP-seq experiments, we determined not only the filtered peaks for each ChIP-seq experiment but also the consensus motif binding sets for each transcription factor. Thus, for each binding site in the genome, the user can see which ChIP-seq experiment has an overlapping peak (technically this means a peak summit position within 50 bp in either direction). In this way, having the three chosen experiments and the consensus motif, we can count how many sites have overlapping peak summits for each of the seven possible combinations. In the VennView, the user can see these values in a Venn diagram. This can be useful for comparing three experiments with not only the same antibody, but from different tissues / developmental stages / treatments and also in examining the extent of overlapping binding of a given transcription factor and its co-factors or co-bound proteins.

In the recent release, the ChIPSummitDB contains data from the analysis of 3727 human ChIP-seq experiments. In the **ExperimentView**, the user can see an overview of the main attributes for each experiment. Most importantly, there are links to the NCBI SRA database (26). The number of sequencing reads and the number of filtered peaks called by the HOMER findPeaks program is also listed (27). During the analysis pipeline, we have determined the most enriched *de novo* and known motifs for each experiment. The link for the results of this search is also located on this page.

The **GenomeView** allows users to see the database content (consensus binding sites, peaks, etc.) in a genomic context through a web browser interface. This view is implemented in the JBrowse framework (34). The GenomeView can be used as a standalone web page where the users can select tracks to load and display. Users can select from the 292 consensus motif sets, the 3727 experiments or from miscellaneous tracks like genomic features or know SNPs. Therefore, users can compare any combinations of experiments versus consensus motif binding sites. Users can also get to this JBrowse interface from the MotifView, the PairShiftView, and the VennView after selecting up to three experiments. In this case, the consensus motif set of the chosen motif and the overlapping peaks and summits of the three chosen experiments will be displayed initially. The user can display any other previously mentioned tracks. Our database provides a comprehensive catalogue of experimentally verified transcription factor binding sites in the human genome. As the cost of whole genome sequencing is drastically decreasing, the number of variations associated with a certain phenotype is rapidly increasing. Most of these variations are in intronic or intergenic regions. Therefore, there is a great interest in determining the overlap of transcription factor binding sites. The **dbSNPView** allows users to check these cases. Users can enter either a genomic region or a dbSNP ID (24). In the first case, the webpage will then display the given region with the variations on it and also the overlapping consensus motif binding sites from our database. The overlapping SNPs are highlighted in red. Either clicking on them or entering the dbSNP accession number directly into the search field leads to the enlarged dbSNPView. Here the reference genome sequence together with the logo of the consensus motif and the overlapping SNPs can be seen. This view is useful for examining how severely the altered base can affect the transcription factor binding. There is also a button to check which experiments give an overlapping peak with the given variation.

### The Novelty of the ChIPSummitDB

The ChIPSummitDB is the first ChIP-seq related database that analyzes and shows the peak summit and consensus motif binding site center distances. The web interface of the database provides several tools for displaying this kind of topological data. In the MotifView, every experiment (antibodies used for the immunoprecipitation) that is above the threshold number of overlapping peaks for a chosen consensus motif is shown. Either direct binding (in the cases where the immunoprecipitated protein recognizes and binds to the given binding site) or indirect binding is indicated. We hypothesize that the standard deviation of the distances of the peak summit and binding site centers shows the directness of the binding. For example, based on the standard deviations, there are at least three groups of interacting proteins at the YY1 consensus motif binding site (Fig. 3a, modified from: http://summit.med.unideb.hu/summitdb/motif_view.php?maxid=10000&minid=1&mnelem=1000&mxelem=120000&motive=YY1). In the first group, the YY1 ChIP-seq experiments have a standard deviation value between 16 and 22 (35). In the next layer, the CTCF and the cohesin proteins and some other TF and co-factor have standard deviation values between 23 and 27 (6). Interestingly, there are many other ChIP-seq experiments, which have a standard deviation above 27 but still have more than 1000 peaks (the threshold value set here), which are overlapping with the YY1 consensus motif binding sites throughout the genome. If we further analyze the scatterplot, we may notice that the YY1 experiments are grouped around 2 at the X-axis, while the CTCF experiments are grouped around −2, as most of the other TFs and cofactors from the third group. Based on these observations, we can hypothesize that the YY1 binds to its cognate binding site. The CTCF-cohesin complex binds to the already bound YY1 protein in the same horizontal arrangement as can be seen at the CTCF binding sites (Fig. 3b, modified from http://summit.med.unideb.hu/summitdb/motif_view.php?maxid=10000&minid=1&mnelem=100&mxelem=120000&motive=CTCF). The proteins in the third group most likely bind to the CTCF-cohesin complex. There are, of course, not only CTCF-cohesin proteins in the second group (e.g. SRF or auts2) (36). Proteins in the second group are most probably directly bound to the YY1 protein.

**Fig. 3.**
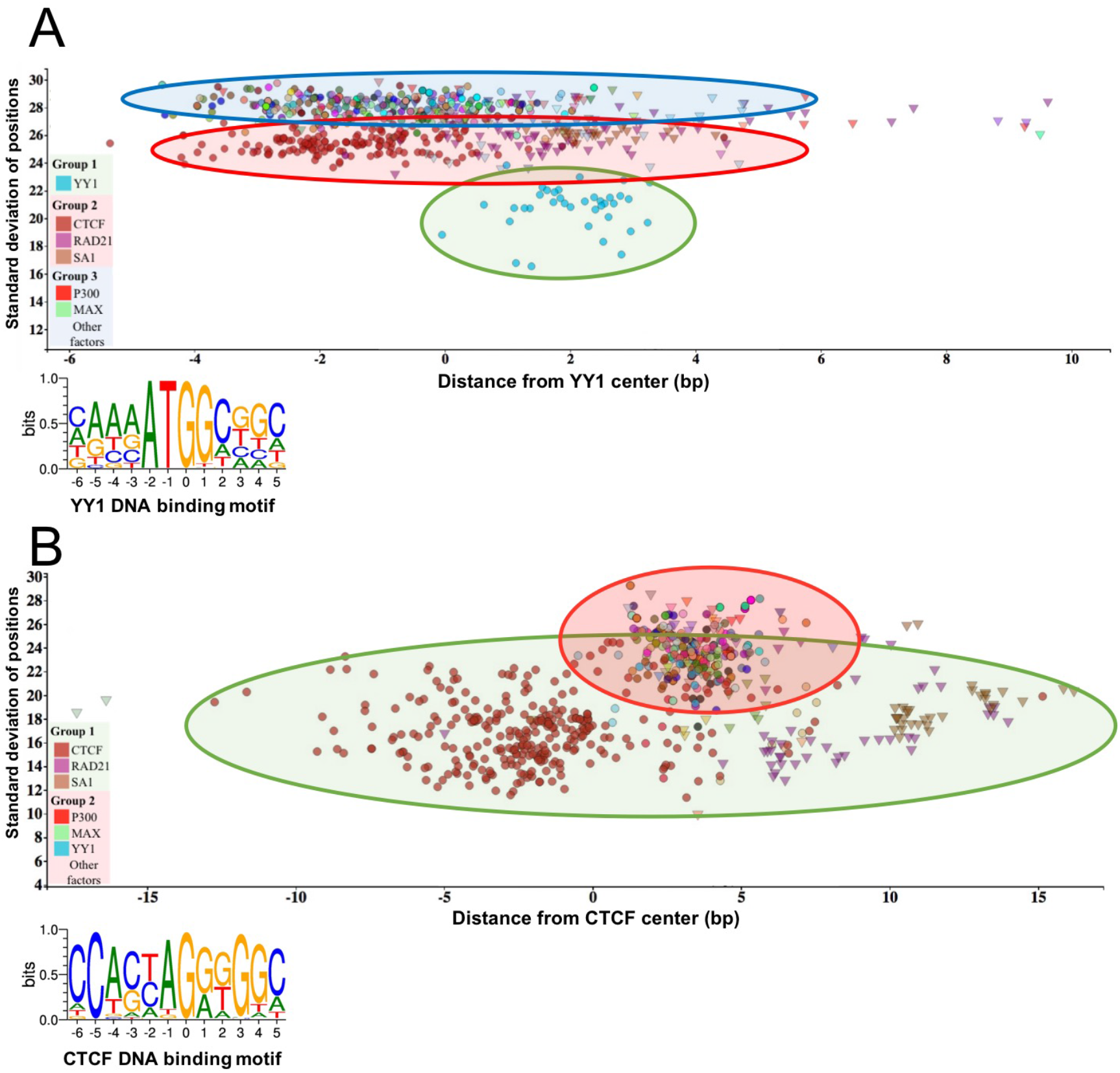
The standard deviation of the distances of the peak summit and binding site centres shows the DNA-protein proximity. Each scatter represents average summit position from a single ChIP-seq experiment. The X axis represents the distance from the binding site centre, which position is marked by „0” in the binding motif logo. The standard deviation of the summit- motif centre distances are shown on the Y axis. A) The proteins, which show interaction with YY1 binding sites, are arranged in three groups. The lowest SD (between 16 and 22) belong to YY1 protein, which binds directly to the YY1 DNA binding motifs. In the second layer, CTCF and cohesin subunit (RAD21, SA1) ChIP-seq signals are the most common. The third group with high SD, above 27, represents a diverse population, which consist of ChIP-seq experiments with different protein targets and more than 1000 overlapping peaks. The P300 and MAX proteins from group 3 are labelled by red and green colours, respectively. The figure was slightly modified and adapted from ChIPSummitDB website: http://summit.med.unideb.hu/summitdb/motif_view.php?maxid=10000&minid=1&mnelem=1000&mxelem=120000&motive=YY1 B) In the case of the CTCF binding sites, only two layers can be distinguished. In the first group the directly interacting CTCF, RAD21 and SA1 proteins can be found, while the YY1, P300, MAX and other proteins are presented in the second group. Please note that the relative position of the YY1 and CTCF proteins to each other is the same on both plots. The figure was slightly modified and adapted from ChIPSummitDB website: http://summit.med.unideb.hu/summitdb/motif_view.php?maxid=10000&minid=1&mnelem=5000&mxelem=120000&motive=CTCF

Besides the already analyzed CTCF-cohesin topology, the ChIPSummitDB provides further evidence that the peak summit versus the binding site based analysis indicates topological arrangements for TF-DNA complexes. One good example is the GATA1∷TAL1 composite element (http://jaspar.genereg.net/matrix/MA0140.2/) (37). Here, the ChIPSummitDB clearly confirms the experimental results. The topological arrangement of the two transcription factors is clearly visible at both the MotifView (Fig. 4a, modified from: http://summit.med.unideb.hu/summitdb/motif_view.php?maxid=2000&minid=1&mnelem=1000&mxelem=120000&motive=GATA1%3A%3ATAL1,withthe“onlydirectbinding”button) and at the PairShiftView (Fig. 4a, modified from: http://summit.med.unideb.hu/summitdb/paired_shift_view.php?exp1=220&exp2=218&exp3=undefined&motive=GATA1::TAL1&motifid=89&limit=25&low_limit=-25&formminid=1&formmaxid=10000&mnelem=100&formmaxelem=120000).

**Fig. 4.**
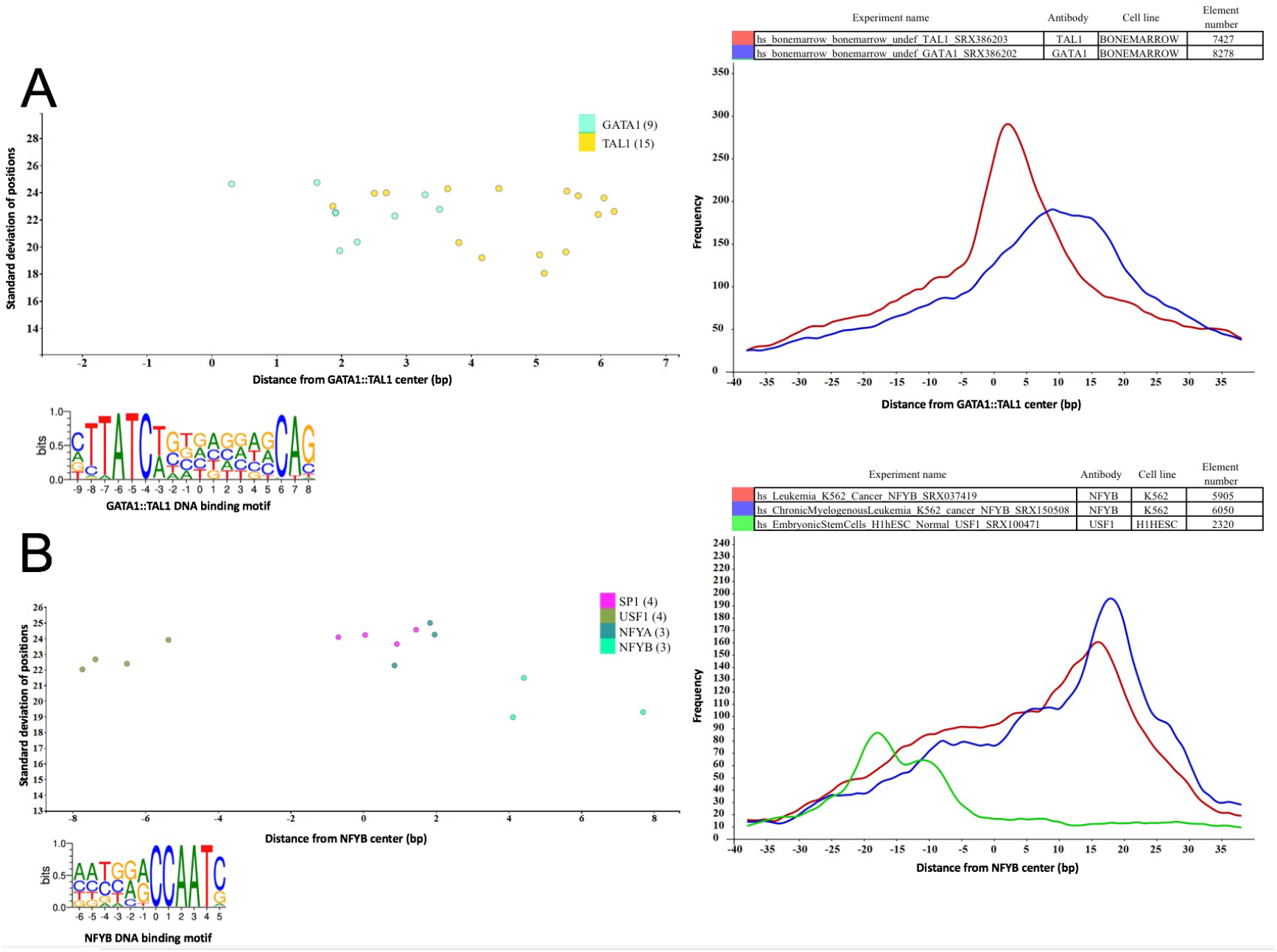
Binding sites based analysis of topological arrangements of TF-DNA complexes as visualized in MotifView and PairShiftView. The plots show the preferred positions of different proteins on A) GATA1∷TAL1 binding sites and on B) NFYB binding sites. The scatterplot follows the same logic as shown on Fig. 3. The figures derive from ChIPSummitDB, although the scatters were filtered to show only the presented factors. GATA1∷TAL1: http://summit.med.unideb.hu/summitdb/motif_view.php?maxid=2000&minid=1&mnelem=1000&mxelem=120000&motive=GATA1%3A%3ATAL1; NFYB: http://summit.med.unideb.hu/summitdb/motif_view.php?maxid=2000&minid=1&mnelem=2000&mxelem=120000&motive=NFYB. The histograms (at right) show the distribution of the summits relative to the midpoint (motif centers). The horizontal axis shows the distance from motif center, measured in base pairs. The vertical axis displays the distance frequency of summits at the given positions. Each ChIP-seq experiment is represented by a frequency curve A) GATA1:TAL1: blue - SRX386203, red - SRX386202; B) NFYB: red - SRX037419, blue -SRX150508, and green - SRX100471), which are smoothed with a rolling mean with a 5 bp window. Element numbers in the tables indicates the number of peak regions obtained in a ChIP-seq experiment, which overlap with a particular consensus motif binding site set. Figures are adapted from ChIPSummitDB website: GATA1∷TAL1: http://summit.med.unideb.hu/summitdb/paired_shift_view.php?exp1=220&exp2=218&exp3=5&motive=GATA1%3A%3ATAL1&motifid=89&limit=40&low_limit=-40&mnelem=100; NFYB: http://summit.med.unideb.hu/summitdb/paired_shift_view.php?exp1=2301&exp2=761&exp3=1597&motive=NFYB&motifid=175&limit=40&low_limit=-40&mnelem=2000

Theoretically, we expect that the peak summit will be in the middle of the transcription factor binding site. We have already shown, in the case of the CTCF-cohesin complex, that this is not necessarily true (38). The ChIPSummitDB provides even more extreme cases. For example, in the case of the NFYB motif observed in the MotifView (Fig. 4b, modified from: http://summit.med.unideb.hu/summitdb/motif_view.php?maxid=10000&minid=1&mnelem=2000&mxelem=120000&motive=NFYB), the average distance values that the NFYB binds are upstream, while the USF1 binds downstream of the motif. If we further scrutinize the topology on the PairShiftView, which shows the real distribution of the distances, we can see even more extremities (Fig. 4b, modified from: http://summit.med.unideb.hu/summitdb/paired_shift_view.php?exp1=2301&exp2=761&exp3=1597&motive=NFYB&motifid=175&limit=40&low_limit=-40&mnelem=2000). In the case of NFYB, most of the distances between the overlapping consensus motifs and peak summits are around the +15 - +17 positions. In the case of USF1, however, the majority of the distance values are clustered around the −18 position. It is also remarkable that in the distribution curves, there are also other smaller shoulders.

## Conclusions

ChIPSummitDB is the first ChIP-seq database based on a transcription factor binding site centered analysis of peak summits. The database convincingly confirms our previous hypothesis that if the different proteins are sitting on the DNA not exactly above each other, then the average peak summit position will display a shifted value relative to the motif center. There can be numerous reasons for this phenomenon. One obvious example is when two different transcription factor binds nearby on a composite element (37). Surprisingly, there are cases, when a transcription factor is bound to its cognate site in the DNA and somehow a different protein without a DNA binding domain that is bound to that TF gets so close to the DNA double helix that it will crosslink to it during the experiment. This will result in a shifted peak summit versus motif center value in our analysis (as can be seen in MotifView and PairShiftView in our database). We showed this shift for cohesion proteins (23), but this shift can also be recognized in other cases.

The detailed analysis of the ChIP-seq summit and motif center positions led us to a new hypothesis: Taking a consensus binding site set (ChIP-seq verified binding sites for a given transcription factor), the closer a given protein is to the DNA, the lower the standard deviation of the distances between overlapping peak summits versus motif center pairs. In other words, if a protein is very close to the DNA double helix, which means in most cases that the protein is bound to the DNA, the resulting ChIP-seq peak summits will more likely be centered in the middle of the DNA region covered by the protein. This recognition can help us better understand how the protein complexes are built on the DNA starting from binding of a transcription factor to its cognate binding site.

Besides these completely new features, the ChIPSummitDB provides a comprehensive, experimental based collection of transcription factor binding sites. The site can be browsed in the GenomeView and we have also developed a dbSNP view that allows users to check whether a given SNP is overlapping a TFBS (24,34). This feature can be useful in determining the consequences of non-coding mutations.

## Supporting information

Summary table of the transcription factor motif, which has identified instances in ChIPSummitDB.

Summary table of the ChIP-seq experiments.

## List of abbreviations

bp: base pair
BWA: Burrows-Wheeler Aligner
ChIP-seq: Chromatin Immunoprecipitation Sequencing
CTCF: CCCTC-binding factor
DB: Database
dbSNP: Single Nucleotide Polymorphism Database
ELF1: E74-like factor 1
ETS1: E26 Oncogene Homolog 1
GATA1: GATA-binding factor 1
GM12878: Lymphoblastoid cell line
HOMER: Hypergeometric Optimization of Motif EnRichment
HTML: HyperText Markup Language
HTTP: HyperText Transfer Protocol
ID: identification number/identifier
NCBI: National Center for Biotechnology Information
NFYB: Nuclear transcription factor Y subunit beta
PHP: Hypertext Preprocessor
PWM: Position weight matrix
RAD21: Double-strand-break repair protein (Scc1, Mcd1)
SA1: Stromal Antigen 1
SMC1/3: Structural maintenance of chromosomes proteins
SNP: Single Nucleotide Polymorphism
SQL: Structured Query Language
SRA: Sequence Read Archive
TAL1: T-cell acute lymphocytic leukemia protein 1
TF: transcription factor
TFBS: transcription factor binding site
USF1: Upstream stimulatory factor 1
XML: Extensible Markup Language
YY1: Yin Yang 1
ZNF143: Zinc Finger Protein 143

## Declarations

### Ethics approval and consent to participate

Since this study carries out exclusively in silico analysis of already published data, it does not need ethics committee approval, as it did not constitute biomedical research. However, reporting is consistent with all ethical requirements.

### Consent for publication

No consent for publication is required.

### Authors’ contributions

EB initiated the project. EB and EC conceived and designed the overall project. EC, JK and OPS carried out the data collection. EB carried out the basic data processing. EC designed and carried out the downstream analysis and the generation of data tables. The website was designed and created by MS and TN and LK. EC, TN and LS performed the statistical analyses and data filtering. EB and EC evaluated the results and wrote the manuscript.

### Availability of data and materials

The datasets supporting the conclusions of this article are included within the article and in Additional file 1-3. The full list of guidelines and a glossary of terms can be found in ChIPSummitDB website (http://summit.med.unideb.hu/summitdb/) and in Additional file 1. The scripts used during the analysis are available at the following website: https://github.com/summitdb.

## Acknowledgments

The authors would like to thank Ferenc Marincs and Karen L. Uray for their critical discussion and technical assistance. Data analyses were performed on the High Performance Computing cluster at the Genomic Medicine and Bioinformatic Core Facility of Department of Biochemistry and Molecular Biology, University of Debrecen.

## Additional file 1.

Supplementary notes and figures.

Figure S1: Schematic representation of the initial data processing. Processing starts with data collection and proper naming. After processing and filtering steps, we get the transcription factor binding sites in bed and bedgraph formats.

Figure S2: Summit prediction. Identification of local maxima within peak regions.

Figure S3: Peak filtering according to shape. (A) Peak with well-defined summit. We filtered peaks depending on the symmetry of their two side (summit positions serves as a symmetry axis) (B), the positions of the 2nd and 3rd quartiles (C), and the symmetry between the read coverage of the two strands (D).

Figure S4: Pairing position weight matrices (PWMs) for processed ChIP-seq experiments. (A) 2758 experiments could be paired to a proper JASPAR motif from the downloaded and processed 3727 ChIP-seq experiments (B). This paired to 338 JASPAR CORE motifs from the 579 (C). The result was a table where the PWMs are paired to their corresponding ChIP-seq experiments (D).

Figure S5: Motif optimization. JASPAR CORE motifs were optimized with the findMotifsGenome program, which used the original PWMs and the merged peak region set of corresponding ChIP-seq experiments (determined in the Motif-ChIP-seq experiment pairing step).

Figure S6: Determining motif locations. To identify the location of motif instances, we combined three different motif finding methods (MAST, HOMER, FIMO). The merged peak region set of corresponding ChIP-seq experiments was used in the identification (Figure 6A). To filter the identified motifs, we used the presented formula in Figure 6B. In the case of overlapping motifs, the motif with the highest Weighted Motif score was selected.

Figure S7: Measuring the distances between motif centers and the surrounding summits. We calculated the concrete distance between motifs and the neighboring summits (measured in base pairs). We took into account all of the possible summits from every experiment.

## Additional file 2.

Summary table of the transcription factor motif, which has identified instances in ChIPSummitDB. The table describes the following basic information for the motifs:

1. column: Name of the motif
2. column: JASPAR record/ID
3. column: motifURL
4. column: Motif length
5. column: SRA antibody name paired to a motif
6. column: used ChIP-seq experiments in consensus motif creation

## Additional file 3.

Summary table of the ChIP-seq experiments. The table describes the following basic information for the experiments:

1. column: name of the experiments
2. column: ftp address of SRA files
3. column: layout of experient (paired/single)
4. column: HTTP address of SRA record of ChIP-seq experiment
5. column: Link to SRA record
6. column: all identified peaks
7. column: filtered peak number
8. column: Transcription factor /cofactor + DNA binding motif
9. column: ExperimentView HTTP address
10. column: Link to ExperimentView

