## Supplementary Material for "ChIPSummitDB: A ChIP-seq-based database of human transcription factor binding sites and the topological arrangements of the proteins bound to them"

*ChIPSummitDB is a collection of processed ChIP-seq data. It provides information about the possible direct or indirect interactions between transcription regulatory proteins and their positions in the genome.*

**Data processing steps:**

Step 1: ChIP-seq data collection from public databases.
Step 2: Processing of collected ChIP-seq data using a custom analysis pipeline. The pipeline includes read mapping, motif enrichment analysis, peak prediction, and making coverage track files (bedgraphs and bigwig). The peak region BED files and coverage file (bedgraph) are essential for the next steps.
Step 3: Splitting peak region and summit prediction. The peak region files and coverage files (bedgraph) from the previous step were used to subdivide ChIP-seq regions into discrete signals and find summit regions.
Step 4: Peak filtering. A custom script was created to filter peak regions, depending on their symmetry and shape. The script utilizes the split peak regions and coverage bedgraph files and results in filtered peak region sets in the BED format.
Step 5: JASPAR CORE motif and ChIP-seq data pairing. We paired the JASPAR motif to their corresponding ChIP-seq experiments. The results were stored in a table.
Step 6: Motif optimization. A motif optimization of JASPAR CORE motifs was conducted. To do this, a merged peak region set was created using the filtered peak regions of the corresponding ChIP-seq experiments (determined in the previous step). All JASPAR motifs were optimized using these merged genomic regions, resulting in optimized motifs.
Step 7: Determining motif locations. We used 3 different programs to find the instances of optimized motifs in the genome. As a result, the genomic locations are in BED format.
Step 8: Summit distance calculation. The centers of identified motifs (step 7) served as reference points in the calculation of motif-protein and protein-protein distance calculations.

The results are stored in a MySQL data table, which is available on the ChIPSummitDB website.

**About ChIPSummitDB:**

The main goal of analyzing ChIP-seq experiments is to identify regions in the genome where we find more sequencing reads (tags) than we would expect to see by chance. These regions are called peak regions due to the appearance of the visualized distribution of mapped tags [1]. The peak's summit (maxima) shows the highest coverage of the region and is known to more-or-less coincide with the center of corresponding DNA elements in the case of transcription factors [2]. These summits correlate with the accurate contact positions of the proteins on the DNA and can be used to determine the topological arrangements of the binding proteins relative to the strand-specific transcription factor binding sites (transcription factor binding motifs (TFBMs)). Earlier we showed [3] that the exact positions of DNA binding proteins on the DNA can be extracted from the ChIP-seq data by identifying the peak summit positions.

Our goal was to create a global database which was based on combining the location of identified transcriptional regulatory elements (TREs) with the positional information of the co-bound regulatory proteins (using publicly available ChIP-seq data, targeting as many proteins as we could). By investigating a global picture of different transcription factors and cofactors, we can identify previously unknown transcriptional regulatory networks. Using the database, we can browse co-bound proteins on TREs and acquire information about their positioning relative to each other and the bound transcription factor motif.

**Processing the data**

Data from 4058 ChIP-seq experiments, covering a wide range of proteins and cell types, were collected from the NCBI SRA and ENCODE databases [4,5]. The naming and automatic download of experiments were performed using a custom script. Processing of the downloaded raw data was carried out using a second in-house developed ChIP-seq analysis pipeline [6] involving mapping [7], peak calling [2,8], tagdirectory creation, and data visualization [8]. The workflow of the procedure is visualized in Figure S1. Following this analysis, the semi-processed data were further analyzed.

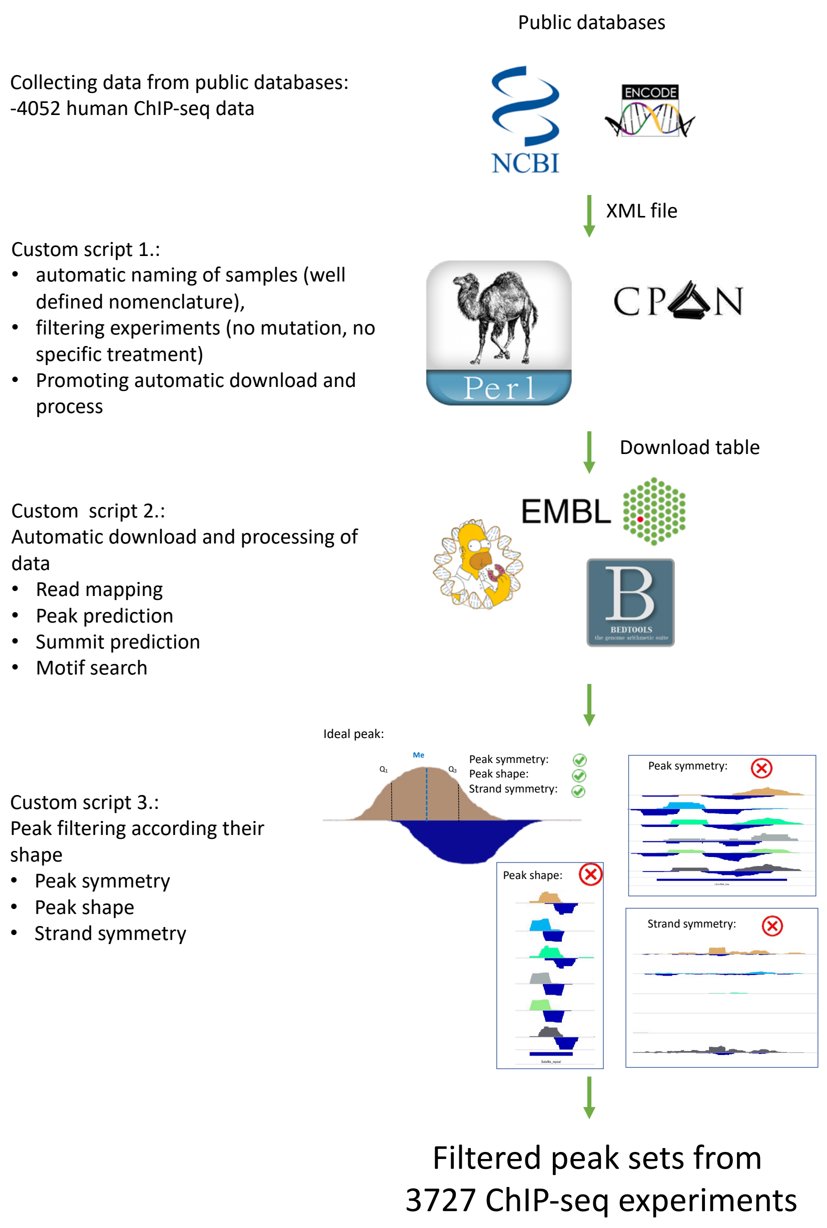


**Figure S1: Schematic representation of the initial data processing.** Processing starts with data collection and proper naming. After processing and filtering steps, we get the transcription factor binding sites in bed and bedgraph formats.

### Peak splitting and summit prediction

We used PeakSplitter, which was developed to split sub-peaks when overlapping peaks are present, for summit predictions. Thus, a more accurate local maxima could be obtained (Figure S2). The peaks for transcription factor binding sequences are usually concentrated to a narrow area, showing a Gaussian distribution due to random fragmentation and their narrow binding surface. This was especially observable after the extension of reads to the expected fragment length [2]. High signal and weak enrichment can indicate insufficient discarding of read duplicates or library preparation artifacts [9].

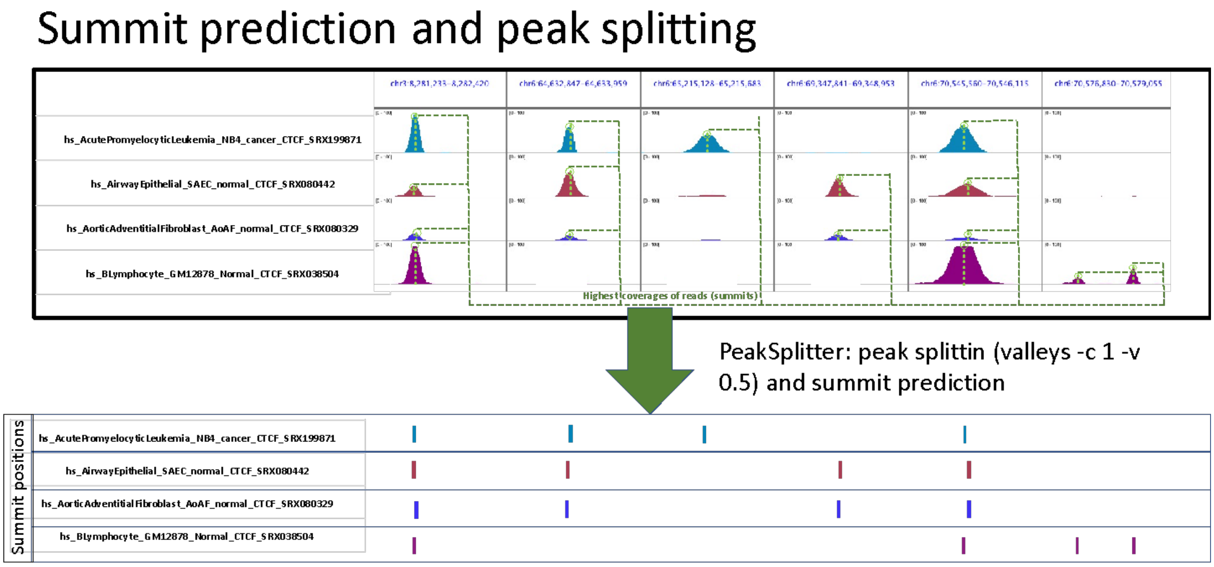


**Figure S2: Summit prediction.** Identification of local maxima within peak regions.

### Peak filtering

Identifying peaks with a well-defined maxima was crucial at the early stage of data processing because false positive peaks could result in false prediction of a protein's position. The peak summits (maxima) show the highest coverage for the peak region and coincide reasonably with the center of the corresponding DNA elements bound by transcription factors. Therefore, the identification of regions suitable for clear determination of the summit position(s) was required. Current software packages use different strategies, such as the evaluation of peak prediction reproducibility or the use of false discovery rates (FDR), for peak prediction [10], which dramatically decrease the false positive rates.

Unfortunately, by using these methods, the filtering algorithms needed to be configured differently for each experiment, which makes the automatization of the processing of large datasets more difficult. For better filtering, we have developed a pipeline, which reduces the false positive discovery rate even further. To avoid false positive results, we filtered out duplicated reads by using a step in the ChIP-seq analysis pipeline and developed a Perl script, which classified and filtered the sub-peaks based on their size and shape.

In these analyses, the peaks are considered coverage histograms and the positions of the median and first and third quartile values were used. The "ideal" transcription factor peak has three attributes: i) the read distribution on both strands have symmetrically curved shoulders, if the median value is the symmetry axis; ii) the peak's shape displays a bell-like curve; iii) the maxima of the ChIP-seq signal is approximately equal between the Watson and the Crick strands (Figure S3a). The first two steps of the filtering analysis are required for filtering out peak positions that have large gaps in their ChIP-seq signal intensity even after the read extension by the peak caller software. The

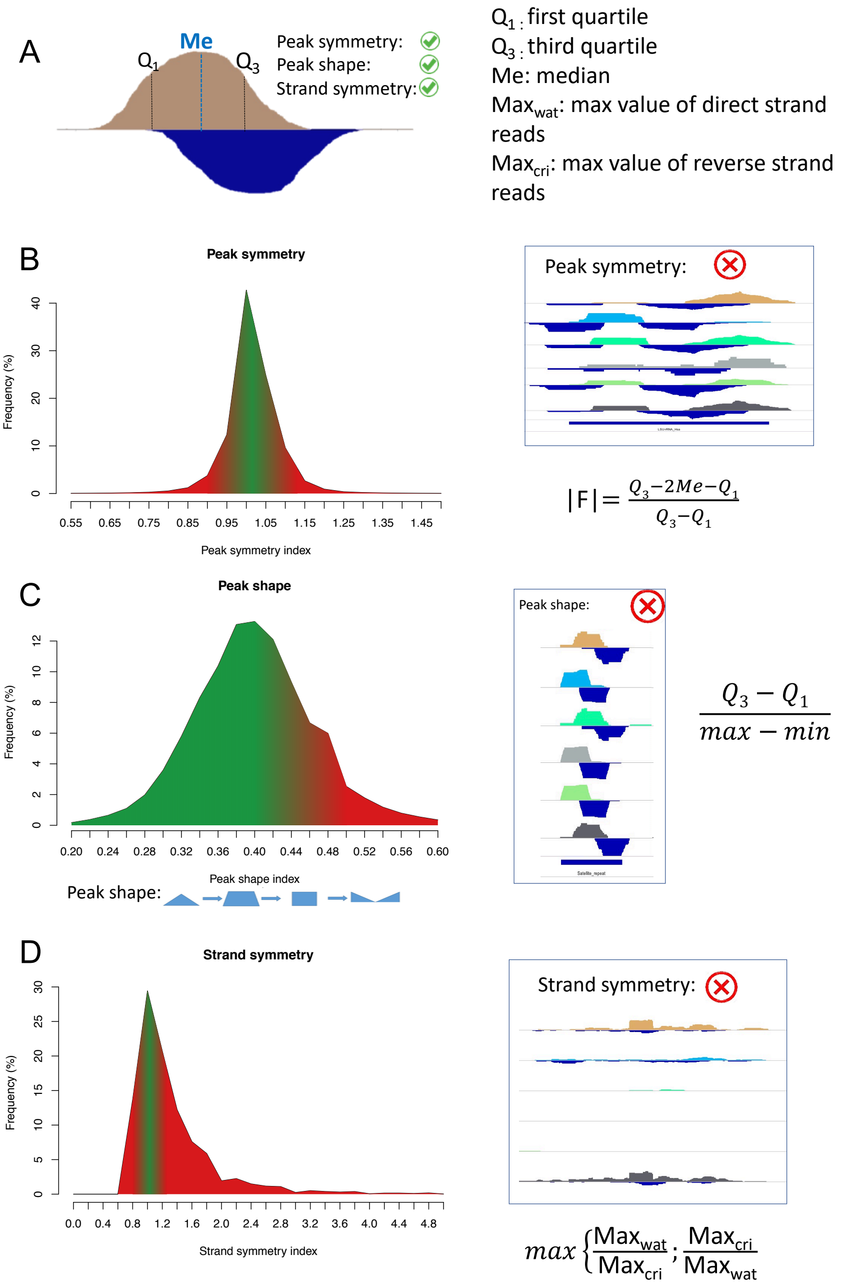


**Figure S3: Peak filtering according to shape.** (A) Peak with well-defined summit. We filtered peaks depending on the symmetry of their two side (summit positions serves as a symmetry axis) (B), the positions of the 2nd and 3rd quartiles (C), and the symmetry between the read coverage of the two strands (D).

formula in Figure S3b calculates how symmetrical the two sides of the peak are. For this calculation, the maxima were used as the axis of symmetry. The second formula (Figure S3c) quantifies the shape of the peak based on the distances between the minimum, maximum, and $2^{nd}$ and $3^{rd}$ quartile values of ChIP-seq signal intensities within the peaks. The resulting value varies between 0 and 1. If we connect the four above-mentioned values with a straight line (where the x axis represents the position of the signal and y axis represents the signal intensity), the peaks which have a "0" shape value would be shaped like a triangle. In contrast, if the value converges to 0.5, the shape of the peak would resemble a square (Figure S3c). Optimally, the forward and reverse tag counts (in a peak) have, approximately, the same size due to the ChIP-seq method. The third formula calculates the symmetry between the reverse and the direct strand tag counts (Figure S3d).

Due to the ChIP-Seq technology, at each protein-DNA binding site, the tags from the forward strands are located on the left-hand side of the binding site and the tags observed from the reverse strand are located on the right-hand side. This is an aspect which is considered and used by several peak-calling (e.g.: macs2) software to extend reads by an average value during peak identification. We used this parameter to filter data. We calculated forward-reverse maxima distances and values which could be found in the 90th percentile passed this filtering step.

**JASPAR CORE motif and ChIP-seq data pairing**

Identification of the exact positions of TF binding sites is the basis of ChIPSummitDB. These motif positions are not only a collection of regulatory regions, but the motif centers were also used as reference points for summit position analysis. Our primary goal was to create consensus binding site sets for as many transcription factors as possible. To do this, we used the JASPAR CORE database, which is a "curated, non-redundant set of profiles, derived from published collections of experimentally defined transcription factor binding sites for eukaryotes" [11] and incorporates 579 non-redundant motifs. We attempted to collect all motifs with ChIP-seq experiments from our collection. Several motifs were lacking NGS data for historical reasons, thus the JASPAR CORE was built to create families of binding profiles for as many structural transcription factor classes as possible. Despite this, we could allocate only 338 motifs to at least one ChIP-seq experiment, because in the available human data, no sequence and NGS data were available for the rest of motifs (Figure S4).

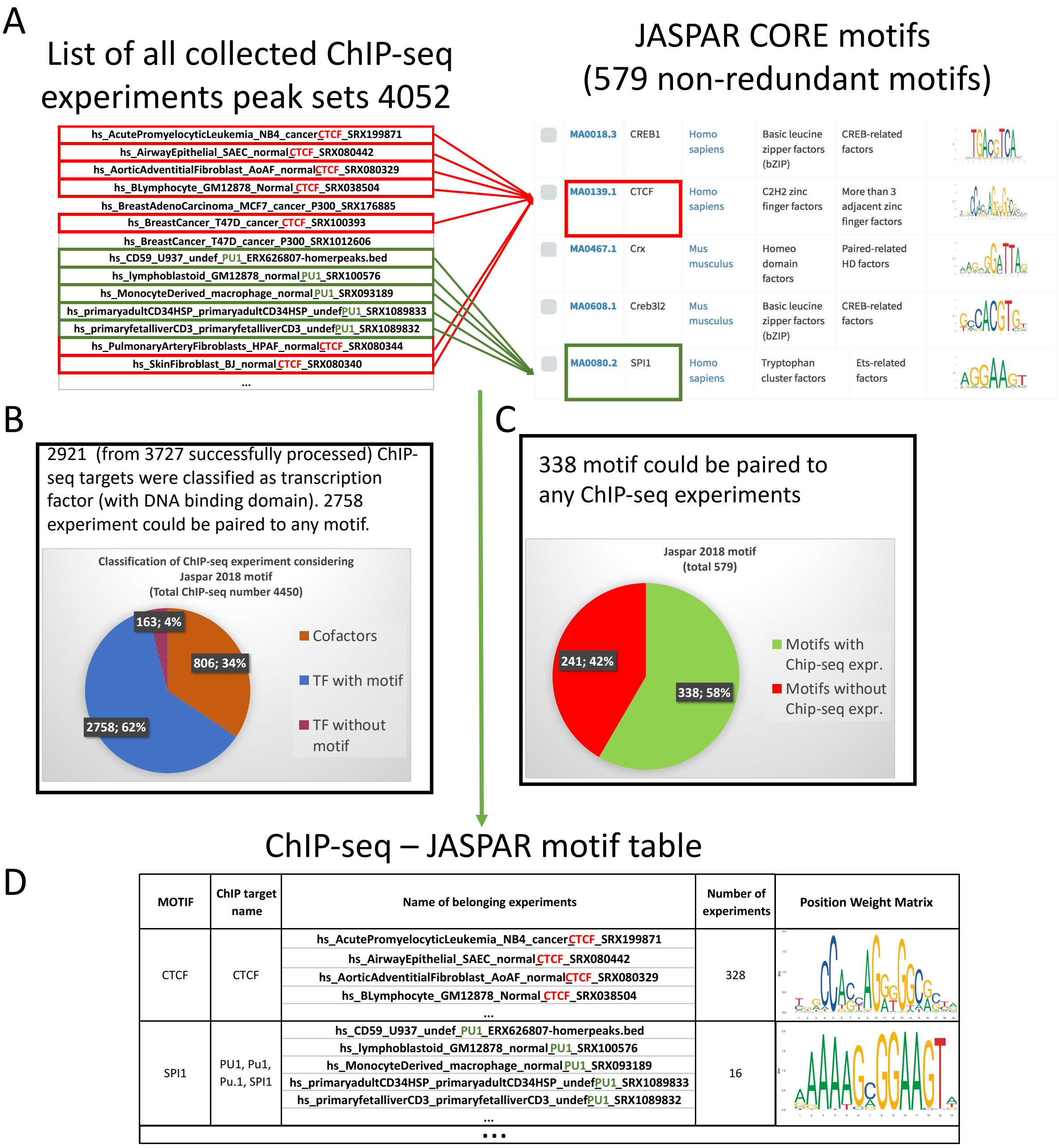


**Figure S4: Pairing position weight matrices (PWMs) for processed ChIP-seq experiments.** (A) 2758 experiments could be paired to a proper JASPAR motif from the downloaded and processed 3727 ChIP-seq experiments (B). This paired to 338 JASPAR CORE motifs from the 579 (C). The result was a table where the PWMs are paired to their corresponding ChIP-seq experiments (D).

### Motif optimization and determining their locations

To optimize the allocated motifs, the peak regions of the corresponding ChIP-seq experiments were scanned for similar motif enrichments [8]. The optimized motifs were manually curated and the most identical ones were paired with the corresponding antibodies (Figure S4). This step maximized the number of specific motif instances, which were identified in the next step (Figure S5).

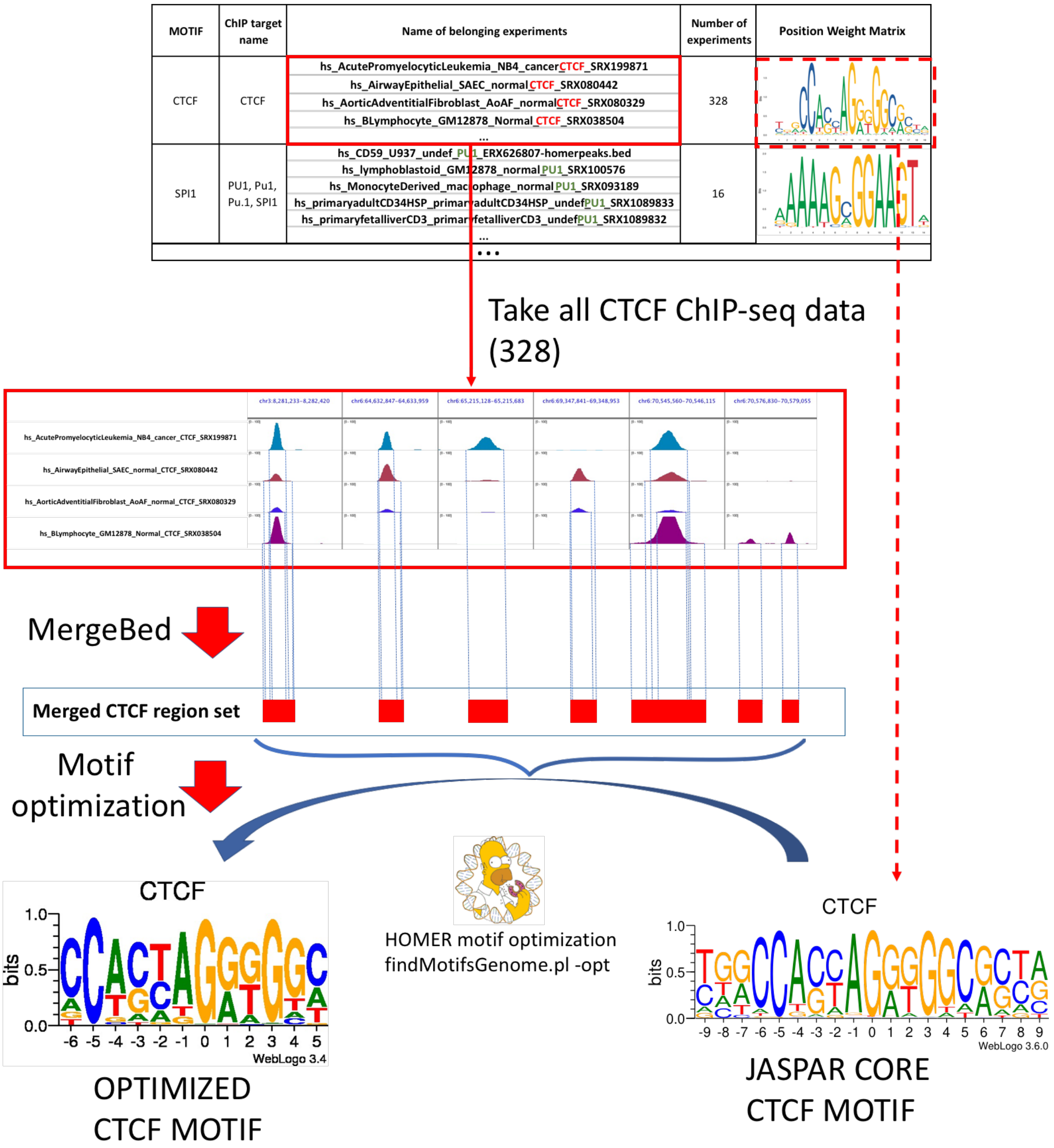


**Figure S5: Motif optimization.** JASPAR CORE motifs were optimized with the findMotifsGenome program, which used the original PWMs and the merged peak region set of corresponding ChIP-seq experiments (determined in the Motif- ChIP-seq experiment pairing step).

Numerous tools can be used to find the occurrences of individual motifs. Instead of choosing one single tool, we combined 3 popular methods: HOMER, FIMO, and MAST [12–14]. The positions which were identified as motifs by at least two programs were filtered in the first step (Figure S6a). Using the default motif scores obtained by the above-mentioned three programs and the distance of the closest summit obtained from the list of paired motif-ChIP-seq experiments, a weighted motif value was calculated. All identified ChIP-seq peaks were coupled with the closest motif possessing the highest weighted motif value. The distance cutoff was +/- 50 base pairs. Following this step, sets of non-redundant motifs were created by filtering out the motifs with identical position and direction (Figure 6A). Even in the case of palindromic sequences, identifying motif directions was possible due to the flanking regions and the positional preferences of the peak summits.

In the previously mentioned step and in the subsequent analysis, closestBed, a tool of bedtools, was used to measure the distance between the center of the motifs and the summits [15]. If the length of an N bp long motif was even, then the (N/2)+1 bp from the 5' end of the sequence was considered as the center of the motif. We created individual summit position pools for all motifs from their respective ChIP-seq experiments. Then, the identified motifs and summits were combined using the closestBed program. This step resulted in a table where all of the summit positions from the proper set are shown together with one or more nearest motif instances. Distances between the centers of the motifs and the summits were calculated this way. Both this distance and the score of the motif were taken into account during the coupling of the most probable motifs with each of the summits. We combined these scores into a formula, and the motif with the calculated highest score was picked for each summit position (one summit could have more than one motif in its vicinity, but only the strongest motif was selected for the following steps). The formula for this calculation can be found in Figure S6b (WMs). The same motif was frequently coupled to summits from different experiments. To avoid redundancy, we removed the duplicates. Thus, we get non-redundant global consensus motif sets for 292 JASPAR CORE matrices. However, 338 motifs could be paired to at least one ChIP-seq experiment, we couldn't find the occurrences of 46 motifs.

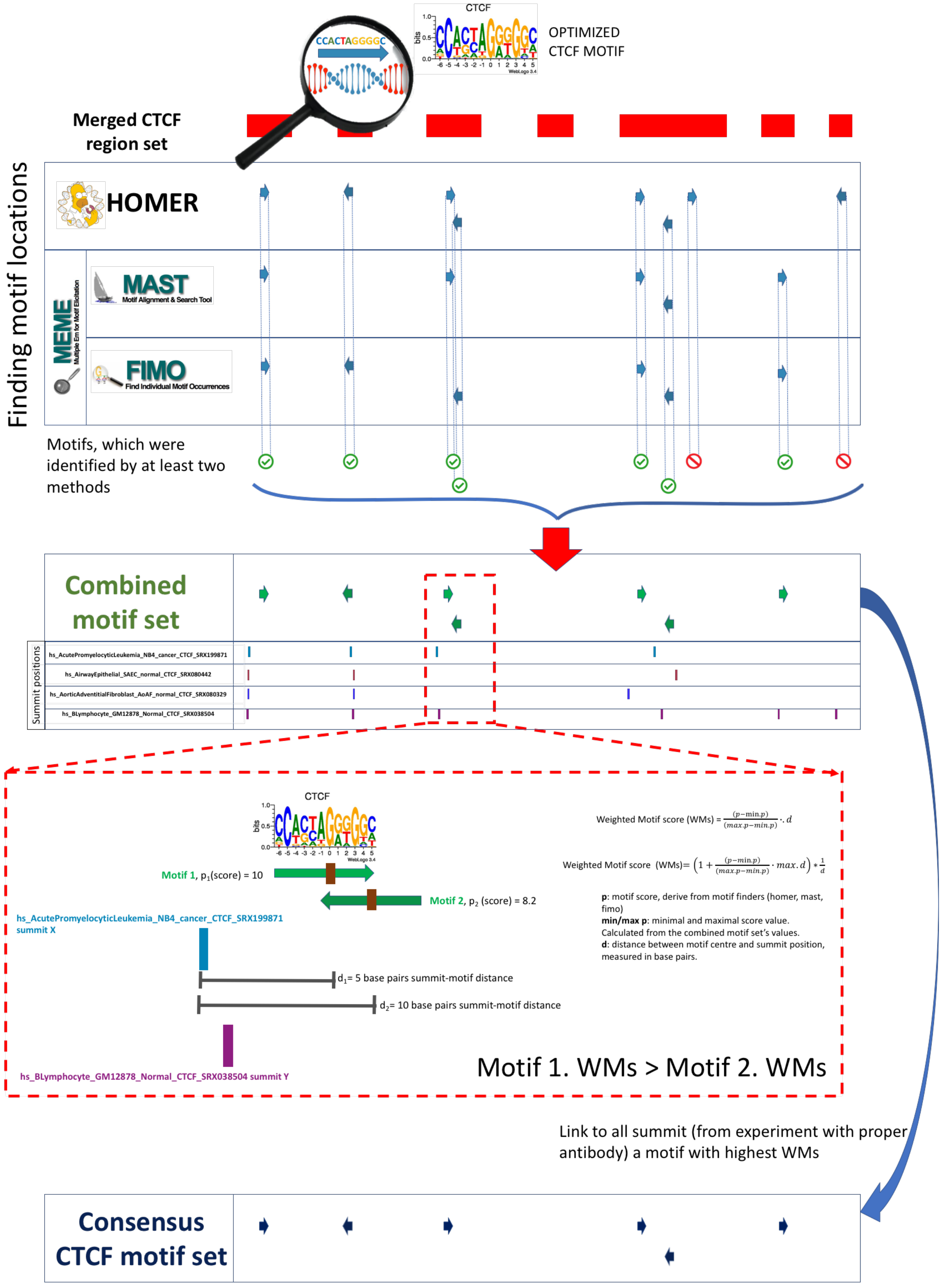


**Figure S6: Determining motif locations.** To identify the location of motif instances, we combined three different motif finding methods (MAST, HOMER, FIMO) [8,12,13]. The merged peak region set of corresponding ChIP-seq experiments was used in the identification (Figure S6a). To filter the identified motifs, we used the presented formula in Figure S6b. In the case of overlapping motifs, the motif with the highest Weighted Motif score was selected.

### Summit distance calculation

The identified consensus sequence locations not only show the genome-wide distribution of transcription binding sites but can be also used as reference points for landscaping of possible co-bindings and the measurement of motif-protein or protein-protein distances. All motif occurrences obtained from every set were screened to identify ChIP-seq experiments containing peak summits in the +/- 50 bp vicinity to the motif center and the distances between motif centers and summit positions were calculated. The resulting distance tables can be examined for either genome-wide or local data. The genome-wide analysis can highlight large-scale information about protein positioning, for example, co-location frequency, location preferences between proteins, possible members of complexes, and patterns in the protein composition of different regulatory regions. In addition to the frequency and the median/average values, both calculated from the measured distances, the standard deviation can also be informative. The preferred position of a particular factor has a larger standard deviation (in relation to the positions of the motif centers) if it is physically far from the reference point (Figure S7).

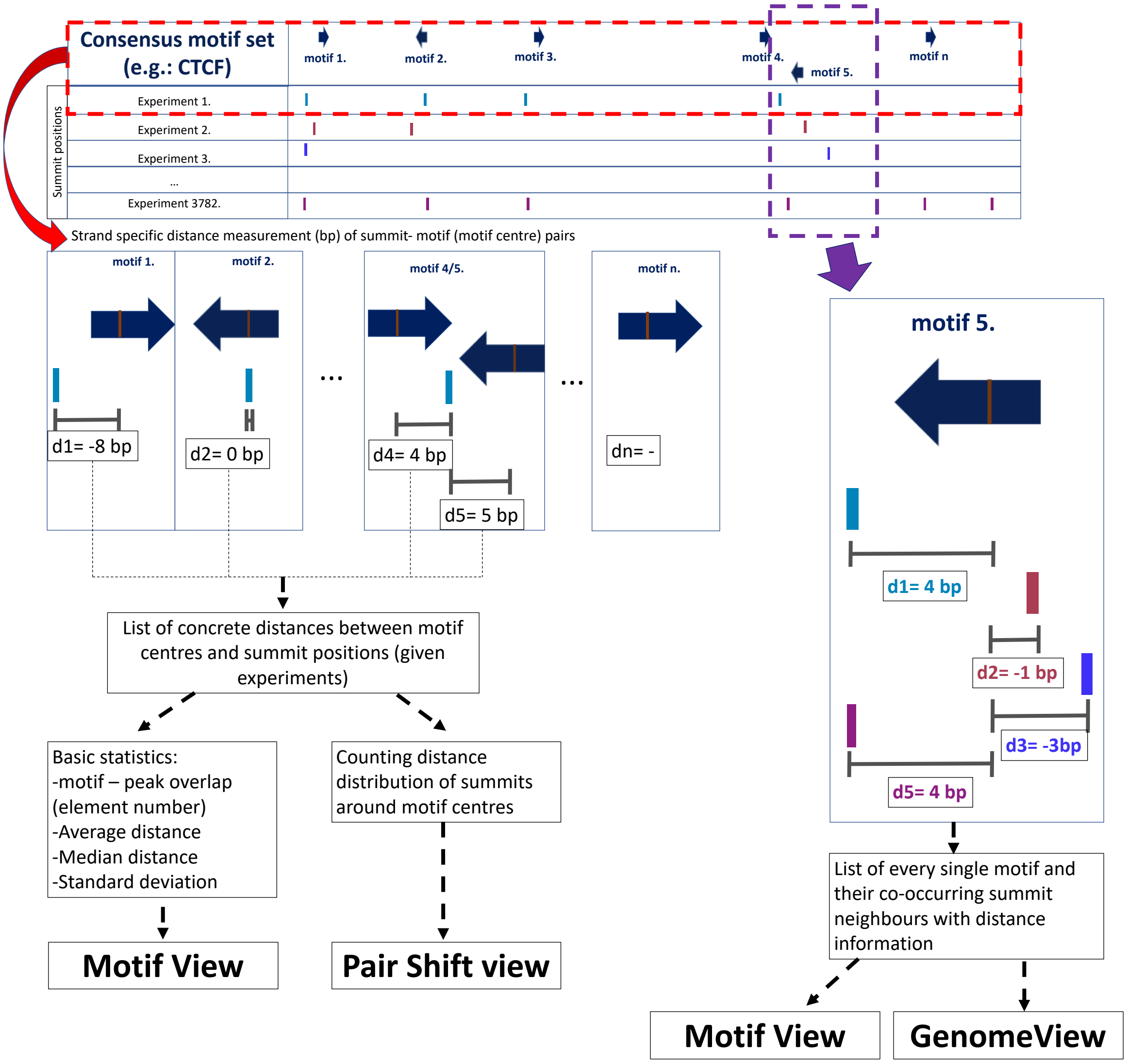


**Figure S7: Measuring the distances between motif centers and the surrounding summits.** We calculated the concrete distance between motifs and the neighboring summits (measured in base pairs). We took into account all of the possible summits from every experiment.

In ChIPSummitDB, the examination of a specific region of the genome is also possible. Examining the summit positions at a specific motif can provide detailed information about the composition of regulatory complexes and their topology, and a comparison of different cell lines is also possible.

**Glossary**

**Average distance**: The average of distances between each summit and motif center pair for a given ChIP-seq experiment and consensus motif pair.

**Consensus binding site:** One of the motifs from the **Consensus motif binding site set**

**Consensus motif binding site set**: The most probable genome-wide binding site set of the transcription factors with a common JASPAR Core motif. A set contains the mapped TFBS matrix genomic positions, which overlap with a corresponding peak region (i.e. the ones with the assigned antibodies). These sets represent all possible experimentally validated binding sites for a given transcription factor.

**ChIP-seq**: A functional genomics technique involving Chromatin immunoprecipitation of a protein-nucleic acid complex followed by next-generation sequencing of the bound nucleic acid. It is used to determine, on a genome-wide basis, the binding position of proteins on chromosomal DNA.

**ChIP-seq experiment**: An experiment using the **ChIP-seq** technique with the following important parameters: the type of the cell and/or tissue, the antibody used for the **IP**, make/model of the sequencing instrument, type of sequencing (single-end or paired-end), sequenced read length, and sequencing depth

**Co-factor**: Here we collect under this name all of the ChIP-seq experiments where the antibody used for the IP is not against a TF

***De novo* motif**: The matrix of an enriched motif determined by the HOMER software. We provide the HOMER determined *de novo* motifs for every ChIP-seq experiment, but they are not used in our pipeline

**Element (number)**: The number of peak regions obtained in a **ChIP-seq experiment**, which overlap with a particular consensus motif binding site set.

**IP**: Immunoprecipitation. An experimental procedure utilizing a specific antibody. It is used for the enrichment of DNA fragments bound by a specific protein.

**JASPAR Core motif set**: JASPAR is a collection of transcription factor DNA-binding preferences, stored as position weight matrices (PWMs). The database contains a non-redundant set of profiles, which are collected from literature data (published and experimentally defined).

**Merged peak set**: Peak regions from different ChIP-seq experiments are merged using bedtools to give every genomic region where there is an experimentally verified binding of a given TF(s).

**Peak**: The pileup of **ChIP-seq** reads on the reference genome. We filter out peaks with non-correct characteristics. We often use the word "peak" to mean "peak region".

**Peak region**: A genomic region, assigned to a chromosome and labeled by the start and end positions, as determined by the peak calling HOMER software, which contains the peak. Ideally, the peak is in the middle of the peak region.

**Peak summit**: The genomic (absolute) or the TFBS-related (as the distance or shift value to the middle of a TFBS) position of the highest (maxima) point of a peak. One peak region can have more than one peak summit. We use PeakSplitter to determine these.

**Shift value (distance)**: The value of the distance between a summit and a TFBS mapped consensus motif center in base pairs. The number can be either negative, positive, or zero.

**Standard deviation (of shift values)**: Here, it is calculated from the shift values between peak summits and the centers of the consensus motif binding sites, which are closer than 50 bp.

**TF**: Transcription Factor. In this database, **TF** is used for any protein which can be immune-precipitated together with its bound DNA, and if a specific binding site from the literature could be assigned.

**TFBS**: Transcription Factor Binding Site. A specific sequence in the genome, which matches with one or more **TFBS matrix**

**TFBS matrix**: A position weight matrix describing the consensus binding site for a **TF**. It is determined using the given JASPAR core matrix for the TF by applying the HOMER motif optimization algorithm on the merged peak region obtained from the corresponding ChIP-seq experiments.

14. Lee MT, Bonneau AR, Takacs CM, Bazzini AA, DiVito KR, Fleming ES, et al. Nanog, Pou5f1 and SoxB1 activate zygotic gene expression during the maternal-to-zygotic transition.

Nature. England; 2013;503:360–4.
15. Quinlan AR, Hall IM. BEDTools: a flexible suite of utilities for comparing genomic features. Bioinformatics. 2010;26.
